# Improving phosphoproteomic kinase activity inference by contextual refinement of Kinase Library relationships

**DOI:** 10.64898/2026.11.04.764166

**Authors:** Yizhi Zhang, Menghuan Zhang

**Author notes:** Correspondence: Menghuan Zhang, 1239 Siping Road, Shanghai 200092, China.

## Abstract

Kinase activity inference from phosphoproteomic data relies on kinase-specific substrate sets defined by kinase–substrate relationship (KSR) priors. Kinase Library (KL) expands candidate substrate coverage using experimentally derived kinase specificity profiles, but its predictions reflect sequence compatibility rather than established intracellular regulation. We benchmarked KL-expanded KSR priors using 222 perturbation phosphoproteomic samples covering 396 annotated activation or inhibition events across 78 kinases. Broad KL expansion increased the number of scoreable kinases from 108 to 391, but reduced perturbed-kinase discrimination, directional accuracy and ranking performance. Higher error rates occurred among kinases with limited curated substrate support, while broad expansion also increased redundancy among inferred activity profiles. To improve the reliability of motif-derived relationships, we constrained KL-derived KSRs using predicted protein co-localization and protein-association evidence. Protein-association filtering consistently improved inference performance, reduced activity redundancy and provided a more favorable accuracy–coverage balance than co-localization filtering. The selected protein-association-filtered prior retained activity estimates for 308 kinases with an area under the receiver operating characteristic curve (AUROC) above 0.75 and extended inference to 200 kinases not scoreable using curated KSRs, which were significantly enriched for dark kinases. These newly scoreable kinases showed structured activity patterns across perturbation contexts and interpretable signaling associations. Our results show that protein-association evidence can refine motif-derived KSR priors while preserving broad kinome coverage, providing a practical framework for more reliable phosphoproteomic kinase activity inference.

## Introduction

Protein kinases are central regulators of cellular signaling and control diverse biological processes through phosphorylation of specific protein substrates. Because kinase abundance does not necessarily reflect catalytic activity, coordinated phosphorylation changes across kinase-specific substrate sets are widely used to infer upstream kinase activity from phosphoproteomic data[1–5]. These substrate-based approaches have been applied to characterize signaling responses, identify dysregulated pathways and prioritize potential therapeutic targets. However, the reliability of kinase activity inference depends strongly on the kinase–substrate relationship (KSR) prior used to define kinase-specific substrate sets. Incomplete or context-inappropriate KSRs can influence inferred activity scores, kinase rankings and even the predicted direction of kinase regulation[6–8].

Experimentally curated KSR resources, including PhosphoSitePlus and PTMsigDB, provide relatively high-confidence evidence of kinase–substrate regulation[3, 9]. However, these resources cover only a limited fraction of experimentally observed phosphosites and are strongly biased toward well-characterized kinases, substrates and signaling pathways[10, 11]. As a result, many kinases, particularly less extensively studied members of the kinome, are represented by few or no quantified substrates in individual phosphoproteomic datasets. This limited coverage restricts the number of kinases for which activity can be inferred and motivates the use of computationally predicted KSRs.

Kinase Library (KL) expands candidate KSR coverage by assigning upstream kinases based on experimentally determined kinase substrate-motif specificity profiles[12, 13]. These predictions provide a systematic description of intrinsic kinase sequence preferences and have enabled large-scale substrate assignment beyond existing curated resources. However, a high KL percentile score indicates that a phosphosite sequence is compatible with the substrate preference of a kinase; it does not establish that the kinase regulates this site in a specific cellular context. Intracellular phosphorylation depends on additional factors, including protein abundance, subcellular organization, docking interactions, protein complexes and stimulus-dependent signaling states[14]. Thus, increasing motif compatibility alone does not necessarily translate into more reliable inference of intracellular kinase regulation. Broad incorporation of motif-compatible relationships may increase substrate coverage while also introducing candidate regulatory links that do not operate in the measured biological context, thereby increasing redundancy among inferred kinase activities.

Recent benchmarking studies have shown that kinase activity inference is sensitive to the evidence composition and confidence of the underlying KSR prior, including differences between curated and predicted KSR resources[6–8]. However, these studies have primarily focused on comparing existing KSR resources rather than developing strategies to refine computationally predicted KSRs. Consequently, how motif-derived predictions can be refined using independent cellular-context evidence remains unclear.

In particular, it remains unclear how KL-based expansion affects the balance between kinase coverage and inference accuracy, whether expanded relationships introduce redundant or less reliable kinase activity estimates, and whether cellular-context evidence can distinguish biologically supported relationships from motif-compatible candidates.

Here, we benchmarked KL-expanded KSR priors using perturbation phosphoproteomic datasets with annotated kinase activation or inhibition events. We evaluated how KL percentile thresholds and per-kinase substrate constraints affected kinase coverage, perturbed-kinase discrimination, directional accuracy, ranking performance and activity redundancy. We further examined whether inference errors were concentrated among specific kinase groups or among kinases that became evaluable only after KL expansion. To improve the reliability of motif-derived relationships, we applied predicted co-localization and protein-association constraints and compared their effects on the accuracy–coverage trade-off and kinase-level activity specificity. Finally, we defined practical operating regimes for different analytical objectives and demonstrated that the selected context-filtered prior enabled biologically interpretable activity inference for newly scoreable kinases, including understudied regions of the kinome.

## Experimental Procedures

### Benchmark phosphoproteomic datasets and perturbation annotations

Benchmark perturbation phosphoproteomic datasets were obtained from the benchmarKIN R package[7]. Phosphorylation changes were represented as log2 fold changes between perturbed and matched control conditions. Perturbation annotations, including target kinases and perturbation direction (activation or inhibition), were also obtained directly from benchmarKIN. Target kinase gene symbols were mapped to reviewed canonical UniProt entries (release 2026_01)[15]. Only samples with annotated human protein kinase targets were retained. The final benchmark set contained 222 samples, including 168 inhibition samples and 54 activation samples.

Each perturbation event represented one annotated target kinase associated with a given treatment. Because a single treatment condition could be associated with multiple target kinases, the 222 samples corresponded to 396 perturbation events involving 78 target kinases. Each event was assigned a perturbation direction according to the benchmarKIN annotation, with inhibition expected to decrease and activation expected to increase target kinase activity. These annotated perturbation events were used as the benchmark reference for evaluating kinase activity inference across different kinase– substrate relationship (KSR) libraries.

### Curated KSR library

The curated KSR library was obtained from benchmarKIN[7]. This collection integrates KSRs from PhosphoSitePlus[9], PTMsigDB[3] after excluding iKiP-DB entries[16], and the GPS 5.0 gold-standard KSR set[17]. Kinase identifiers were harmonized by mapping gene symbols to reviewed canonical UniProt entries. Only relationships whose upstream kinase corresponded to a human protein kinase were retained.

After filtering and identifier harmonization, the curated library contained 313 protein kinases and 4,904 KSRs. The curated KSR set was used as the baseline prior for kinase activity inference and was retained unchanged in all KL-expanded, NetworKIN-expanded and context-filtered priors.

### Construction of KL-expanded KSR libraries

KL-expanded KSRs were generated using the Kinase Library Python package[12, 13]. The phosphosites used for Kinase Library prediction were the unique sites quantified across the 222 benchmarKIN samples. Phosphosites were mapped to reviewed canonical UniProt entries and retained only when both residue identity and position matched the corresponding UniProt sequence. For each retained site, a phosphoacceptor-centered ±7 amino acid sequence window was extracted and used as input to Kinase Library. Kinase Library predictions were generated once for the unified phosphosite set and subsequently used to construct all KL-expanded priors. Candidate upstream kinases with percentile scores ≥90 were retained as KL-derived KSRs. KL-derived KSRs already present in the curated library were removed to avoid duplication, resulting in 1,874,361 KL-derived KSRs involving 388 protein kinases.

A series of KL-expanded priors was generated by combining curated KSRs with KL-derived KSRs selected using different percentile-score and ranking thresholds. KL90, KL95, KL97.5 and KL99 retained KL-derived KSRs with percentile scores ≥90, ≥95, ≥97.5 and ≥99, respectively. For the restricted KL99-T10, KL99-T20 and KL99-T50 libraries, KL-derived KSRs were first required to have percentile scores of at least 99, and only the top 10, 20 or 50 substrate sites by percentile score were retained for each kinase. In all KL-expanded priors, curated KSRs were retained unchanged and KL-derived KSRs were added as candidate motif-compatible relationships.

### Cellular-context filtering of KL-derived KSRs

Cellular-context filters were applied to KL-derived KSRs to prioritize motif-compatible relationships supported by predicted protein co-localization or protein-association evidence. Curated KSRs were retained unchanged in all filtered priors. Each filtering strategy was applied independently to the same set of KL-expanded priors.

### DeepLoc co-localization filtering

Protein subcellular localization was predicted using DeepLoc 2.1[18]. Full-length canonical protein sequences were used as input, and localization predictions were obtained for nine compartments: nucleus, cytoplasm, extracellular space, mitochondrion, cell membrane, endoplasmic reticulum, Golgi apparatus, lysosome/vacuole and peroxisome. A KL-derived KSR was retained if the upstream kinase and substrate shared at least one predicted localization compartment.

### STRING and kinase-interaction network filtering

Protein-association evidence was obtained from STRING (v12.0)[19]. The STRING combined score was used as a confidence measure for functional association between each upstream kinase and substrate. STRING400 and STRING700 retained KL-derived KSRs with STRING combined scores ≥400 or ≥700, respectively.

To complement STRING with kinase-focused interaction evidence, kinase-protein links reported by Buljan et al.[20] were incorporated as additional protein-association support. For each protein-association filter, a KL-derived KSR was retained if the kinase–substrate pair either met the corresponding STRING score threshold or was supported by a kinase–protein interaction in the Buljan et al. network. KSRs lacking support from either source were removed. Representative kinase–substrate networks were visualized using Cytoscape[21].

### STRING evidence-channel decomposition

To assess the contribution of different types of STRING evidence, we decomposed the STRING evidence channels. Two evidence subsets were defined: experimental plus co-expression evidence, and text-mining plus database evidence. Subset-specific combined scores were recalculated from the corresponding channel scores using the STRING probabilistic score-combination procedure. Channel scores were first rescaled from 0–1,000 to 0–1. For each included channel *i*, with rescaled channel score *s_i_*, the STRING prior probability (*p* = 0.041)[19] was removed as

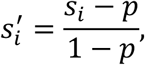

Channels lacking evidence were assigned 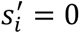.

The prior-corrected channel scores were then combined as

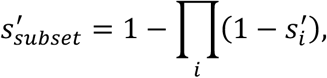

and the prior was added back once to obtain the final subset score,

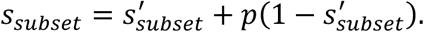

The resulting scores were multiplied by 1,000 and used as subset-specific combined scores.

For comparison, six KL95-based priors were evaluated: unfiltered; STRING400, retaining KSRs supported by either a STRING combined score ≥400 or the Buljan network; Buljan-only, retaining only KSRs supported by the Buljan network; STRING-only, retaining KSRs with a STRING combined score ≥400; experimental + co-expression, retaining KSRs with a corresponding subset score ≥400; and text-mining + database, retaining KSRs with a corresponding subset score ≥400.

### Construction and filtering of the NetworKIN-expanded library

As a complementary analysis, a NetworKIN-expanded library was constructed to evaluate whether cellular-context filtering also improved KSR predictions that already incorporate motif and network information. NetworKIN predictions[22] were obtained from the benchmarKIN data collection deposited on Zenodo, using the file networkin_human_predictions_3.1.tsv. NetworKIN-derived KSRs with scores ≥5 were retained, and predictions already present in the curated KSR set were removed to avoid duplication. The remaining NetworKIN-derived KSRs were added to the curated KSRs to generate the NetworKIN-expanded library, which contained 187 protein kinases and 22,114 KSRs. The same cellular-context filtering framework used for KL-derived KSRs was then applied to NetworKIN-derived KSRs.

### Weighted z-score kinase activity inference

Kinase activities were inferred using a weighted z-score formulation based on the RoKAI approach[4]. Each KSR library was represented as a weighted kinase–substrate network before activity inference. Curated KSRs and NetworKIN-derived KSRs were assigned a weight of 1, whereas KL-derived KSRs were weighted by their Kinase Library percentile score divided by 100.

For each perturbation sample, non-missing phosphosite log2 fold changes were extracted from the phosphoproteomic matrix. The KSR library was restricted to substrate sites quantified in that sample, and kinases with fewer than five quantified substrate sites were excluded from activity inference for that sample. For each remaining kinase *k* in perturbation sample *c*, the activity z-score was calculated as

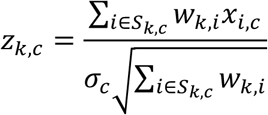

where *x_i_*_,*c*_ denotes the log2 fold change of substrate *i* in perturbation sample *c*, *S_k_*_,*c*_ denotes the set of quantified substrates linked to kinase *k* in that sample, *w_k_*_,*i*_ denotes the weight assigned to the KSR between kinase *k* and substrate *i*, and *σ_c_* denotes the standard deviation of log2 fold changes across all quantified phosphosites in sample *c*. When all KSR weights were equal to 1, this formulation reduced to the standard RoKAI z-score. Positive z-scores indicated increased inferred kinase activity, whereas negative z-scores indicated decreased inferred kinase activity.

### Evaluation design

Performance was evaluated using annotated perturbation events as the benchmark reference. For each KSR library, kinase activity inference produced a sample-by-kinase activity matrix, from which several levels of coverage were defined. Scoreable kinases were defined as kinases with an inferred activity score in at least one sample. Evaluable kinases were defined as annotated target kinases for which activity could be inferred under a given KSR library. Evaluable perturbation events were defined as perturbation events for which the annotated target kinase had an inferred activity score under the corresponding KSR library.

For discrimination and ranking analyses, inferred kinase activity scores were first signed according to the expected perturbation direction. Activity scores were multiplied by +1 for activating perturbations and by −1 for inhibitory perturbations, so that higher signed scores consistently indicated stronger agreement with the expected target-kinase response. Area under the receiver operating characteristic curve (AUROC) was calculated by pooling signed activity scores across all samples. For each sample, all annotated target kinases with available activity scores were added to the positive pool, whereas all other kinases with available activity scores in the same sample were added to the negative pool. Because the number of pooled negative scores exceeded the number of pooled positive scores, a balanced AUROC was computed by randomly sampling negative scores to match the number of positive scores for 1,000 iterations. This metric estimates whether annotated target kinases receive higher signed activity scores than non-target kinases across the benchmark set.

Rank-based metrics were calculated within each sample from signed activity scores. Kinases were ranked in descending order of signed activity score, and the rank of each annotated target kinase was recorded. Within each sample, ranks for all kinases with inferred activity scores were converted to percentile ranks as *r*/*n*, where *r* is the rank of a kinase and *n* is the total number of kinases with inferred activity scores in that sample. The percentile ranks of annotated target kinases were then extracted for evaluation, with lower percentile ranks indicating better recovery. PHit@k% was calculated from the target-kinase percentile ranks. For target kinases evaluated in multiple samples, percentile ranks were first averaged across samples. PHit@k% was then defined as the fraction of evaluable target kinases whose averaged percentile rank fell within the top k% of kinases with inferred activity scores, with k = 10, 15 or 20. Recall@k% was calculated at the sample level. For each sample, the top k% of kinases ranked by signed activity score was selected, and recall was defined as the fraction of evaluable target kinases in that sample that were recovered within this top-ranked set. The reported Recall@k% corresponds to the mean recall across all samples.

Directional accuracy was used to evaluate whether inferred activity changes matched the expected direction of perturbation. For activating perturbations, a positive inferred activity score was considered directionally correct, whereas for inhibitory perturbations, a negative inferred activity score was considered directionally correct. Directional accuracy was defined as the fraction of evaluable perturbation events with the expected activity direction.

### Kinase-level directional error analysis

Kinase-level directional error rates were calculated to assess whether inference errors were concentrated in specific target kinases. For each annotated target kinase, all evaluable perturbation events targeting that kinase were collected under each KSR library. An event was considered directionally mismatched when the inferred activity was negative for an activating perturbation or positive for an inhibitory perturbation. The kinase-level error rate was then defined as the number of directionally mismatched events divided by the total number of evaluable events for that target kinase.

Target kinases were further stratified according to their evaluability under the curated KSR library. Evaluable kinases with curated KSRs were defined as annotated target kinases that were evaluable using the curated library, whereas newly evaluable kinases were defined as annotated target kinases that were not evaluable using the curated library but became evaluable after KL-derived KSR expansion. Kinase group annotations were obtained from KinHub[23] and used to summarize directional error rates across kinase groups.

### Coverage-adjusted utility score

To jointly evaluate discrimination performance and benchmark-event coverage, we calculated a coverage-adjusted utility score for each KSR library:

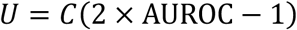

where *C* denotes the fraction of the 396 benchmark perturbation events that were evaluable under the given KSR prior. The term 2 × AUROC − 1 rescales AUROC relative to the random expectation of 0.5, such that random discrimination corresponds to zero utility and higher values indicate better-than-random discrimination. For each context-filtered KSR library, the change in *U* was calculated relative to the corresponding unfiltered KSR library:

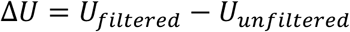

Positive Δ*U* values indicate an improved accuracy–coverage trade-off after context filtering.

### Kinase activity redundancy and distinguishable activity profiles

To assess redundancy among inferred kinase activity profiles, pairwise Spearman’s rank correlation coefficients (*ρ*) were calculated across the benchmark samples for each KSR library. Kinase pairs were included when activity scores were available in at least 10 shared samples. Kinase family annotations were obtained from KinHub[23], and kinase pairs were classified as within-family or between-family.

To estimate the effective number of distinguishable kinase activity profiles, we performed hierarchical clustering on the pairwise activity correlation matrices using 1 − *ρ* as the correlation distance and complete linkage, such that cluster merging was determined by the least similar kinase pair between clusters. Activity clusters were defined by cutting the dendrogram at a threshold of *ρ* ≥ 0.7. Kinases with insufficient observations for correlation analysis were conservatively retained as singleton activity clusters.

To examine whether similarity between inferred kinase activities was associated with overlap in their underlying substrate sets, KSR overlap was calculated for every kinase pair using Jaccard similarity. The association between pairwise KSR Jaccard similarity and the corresponding pairwise activity correlation was quantified using Spearman *ρ*.

### Identification and annotation of newly scoreable kinases

Downstream biological analyses focused on the KL95–STRING400 prior as a representative context-filtered KL-expanded library with broad kinase coverage. Kinases were defined as newly scoreable if an activity score could be inferred using KL95–STRING400 in at least one benchmark sample but not using the curated KSR library. Kinases scoreable using the curated KSR library were used as the reference group where applicable. To assess whether expanded coverage preferentially included understudied kinases, kinases were classified as dark or non-dark using Illuminating the Druggable Genome (IDG) annotations[10, 24]. Dark-kinase proportions were compared between newly scoreable kinases and kinases scoreable using the curated KSR prior using Fisher’s exact test.

### Perturbation-class activity analysis of newly scoreable kinases

To examine whether the inferred activities of newly scoreable kinases differed systematically across perturbation contexts, the 222 benchmark samples were further grouped according to the mechanism of the applied treatment or stimulus. Perturbation classes were designated as focal perturbation classes if they included at least five samples, two independent studies and two distinct perturbing agents. Ten classes met these criteria: RTK stimulation, TCR stimulation, RTK inhibition, PI3K–AKT–mTOR inhibition, RAF–MEK–ERK inhibition, SRC–ABL inhibition, cell-cycle kinase inhibition, DDR kinase inhibition, PKC inhibition and ROCK inhibition.

For each newly scoreable kinase, inferred activities in each focal perturbation class were compared with those in all other samples with the same perturbation direction, which constituted the direction-matched background. Comparisons required at least five non-missing activity measurements in the focal class and 10 in the background. Differences were assessed using two-sided Brunner–Munzel tests, and effect sizes were quantified using Cliff’s delta, calculated as

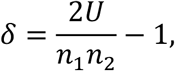

where *U* is the Mann–Whitney statistic and *n*_1_ and *n*_2_ are the sample sizes of the focal perturbation class and direction-matched background, respectively. Positive values indicate higher inferred kinase activity in the focal class than in the background, whereas negative values indicate lower activity.

### Functional characterization of newly scoreable kinases

Functional associations of newly scoreable kinases were evaluated by correlating KL95–STRING400-derived activity profiles with curated-prior activity profiles of reference kinases. For each newly scoreable kinase, Spearman *ρ* was calculated against each kinase scoreable using the curated KSR prior. Kinase pairs with fewer than 10 shared samples were excluded. To quantify the representation of closely related kinases in the reference set, we calculated the proportion of reference kinases belonging to the same kinase family as each newly scoreable kinase.

Gene-set enrichment analysis (GSEA)[25] was performed using GSEApy (v1.1.13)[26] with human Reactome pathways from version 97[27]. Pathway gene sets were restricted to kinases scoreable using the curated KSR prior, and pathways containing 5–31 reference kinases were retained after removing exact duplicate gene sets. For each newly scoreable kinase with at least 10 valid correlations, reference kinases were ranked by Spearman *ρ* and analyzed using preranked GSEA. Pathways with FDR q < 0.05 were considered significant.

### Statistical analysis

Statistical analyses were performed in Python using SciPy (v1.14.1)[28]. Associations between continuous variables were assessed using Spearman’s rank correlation tests. For analyses relating median AUROC to the numbers of scoreable kinases, evaluable kinases, or evaluable perturbation events, ordinary least-squares regression was additionally used to describe linear trends. Regression slopes were rescaled to represent the change in median AUROC per 100 scoreable kinases, per 10 evaluable kinases, or per 100 evaluable perturbation events, respectively.

Differences in phosphorylation changes between STRING-supported and unsupported substrate groups were evaluated using one-sided Mann–Whitney U tests. Enrichment of IDG-defined dark kinases among newly scoreable kinases relative to kinases scoreable using the curated KSR prior was evaluated using a one-sided Fisher’s exact test. Perturbation-class activity differences were assessed using two-sided Brunner– Munzel tests, with Cliff’s delta used to quantify effect size. Exact sample sizes and *P* values are reported in the corresponding figures or figure legends.

## Results

### Broad expansion of motif-derived kinase–substrate relationships increases coverage but reduces inference reliability

We first investigated how expanding kinase–substrate priors with sequence-derived relationships affects phosphoproteomic kinase activity inference. We used Kinase Library (KL)-derived KSRs as a representative motif-based prediction system. The benchmark included 222 perturbation phosphoproteomic samples with annotated kinase activation or inhibition events. For each sample, kinase activity was inferred using either the experimentally curated KSR prior or KL-expanded priors containing increasing numbers of predicted KSRs. Four expanded libraries were generated using KL percentile thresholds (KL90, KL95, KL97.5 and KL99). Three additional libraries retained only the top 10, 20 or 50 KL-derived substrates per kinase from the highest-confidence KL predictions. Activity scores were calculated using a z-score-based approach and evaluated using discrimination, ranking, directional accuracy and coverage metrics (Fig. 1a; see Experimental Procedures).

**Figure 1.**
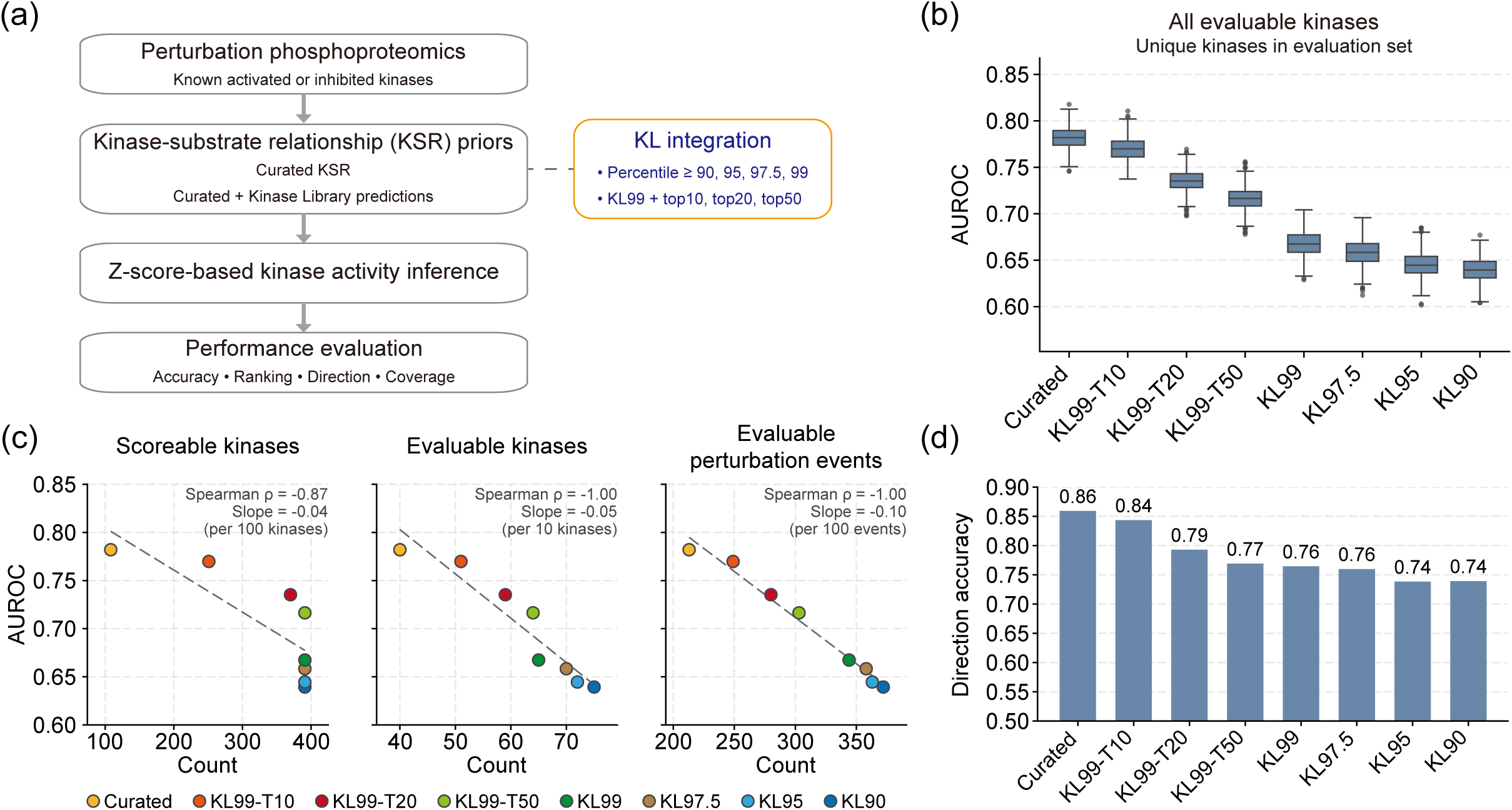
Broad expansion of motif-derived kinase–substrate relationships increases coverage but reduces activity inference accuracy. **a,** Overview of the benchmarking workflow. The benchmark comprised 222 perturbation phosphoproteomic samples, including 168 inhibition and 54 activation samples, corresponding to 396 perturbation events involving 78 kinases annotated as perturbation targets. Kinase activity inference was evaluated using the curated kinase– substrate relationship (KSR) library or Kinase Library (KL)-expanded KSR libraries generated by adding KL-derived KSRs to the curated library. Kinase activities were inferred using a weighted z-score formulation based on the RoKAI approach and evaluated by discrimination, rank-based target-kinase recovery, directional accuracy and coverage. KL99-T10, KL99-T20 and KL99-T50 retained KL-derived KSRs with percentile scores ≥99 and the top 10, 20 or 50 substrates per kinase, respectively. KL99, KL97.5, KL95 and KL90 retained KL-derived KSRs with percentile scores ≥99, 97.5, 95 and 90, respectively. **b,** Area under the receiver operating characteristic curve (AUROC) for distinguishing annotated perturbed kinases from non-target kinases among all kinases with inferred activity across the curated and KL-expanded priors. Box plots show the distributions of balanced AUROC values obtained from 1,000 iterations in which non-target kinase scores were randomly sampled to match the number of annotated perturbed-kinase scores. **c,** Relationship between AUROC and coverage across KSR libraries. Coverage was summarized as the number of scoreable kinases (left), evaluable kinases (middle) and evaluable perturbation events (right). Spearman’s rank correlation coefficients (*ρ*) and fitted slopes are shown. **d,** Directional accuracy across KSR libraries, defined as the fraction of evaluable perturbation events for which the inferred activity change of the annotated target kinase matched the expected perturbation direction.

Despite the substantial increase in kinase–substrate coverage, broader expansion did not improve inference accuracy. The curated prior achieved the highest median AUROC, approximately 0.78. KL99-T10 showed similar performance, whereas AUROC decreased as more KL-derived relationships were incorporated. The broad percentile-based libraries showed the lowest performance, with median AUROC values of approximately 0.64–0.65 for KL95 and KL90 (Fig. 1b). Thus, motif-based expansion alone increased substrate coverage but did not improve recovery of experimentally perturbed kinases.

The reduction in performance was accompanied by a substantial increase in kinase coverage. The curated prior supported activity estimates for 108 kinases, whereas the most permissive KL-expanded prior supported activity estimates for 391 kinases. The numbers of evaluable kinases and perturbation events also increased after KL expansion. However, higher coverage was associated with lower AUROC across expanded priors. AUROC was negatively correlated with the number of scoreable kinases (Spearman *ρ* = −0.87), evaluable kinases (*ρ* = −1.00), and evaluable perturbation events (*ρ* = −1.00) (Fig. 1c).

Directional accuracy showed the same trend. The curated prior recovered the expected activation or inhibition direction in 86% of evaluable events. KL99-T10 retained an accuracy of 84%, whereas accuracy decreased to 79% for KL99-T20 and 74–77% for KL99-T50 and percentile-based libraries (Fig. 1d). Similar trends were observed when activation and inhibition events were analyzed separately, although activation events showed greater variability because fewer samples were available (Supplementary Fig. S1a).

Rank-based analyses further supported these observations. Target-kinase percentile ranks became less favorable as KL thresholds became less stringent. PHit@k% decreased across broader KL libraries, while Recall@k% showed a smaller decline (Supplementary Fig. S1b–d). The AUROC differences between priors remained when analyses were restricted to kinases evaluable with curated KSRs (Supplementary Fig. S1e). AUROC was also positively associated with the fraction of curated relationships retained in each prior (Spearman’s rank correlation coefficient, *ρ* = 1.00; Supplementary Fig. S1f). These findings indicate that broad KL expansion increases kinase coverage but introduces a trade-off with kinase activity inference performance. Complete performance metrics for curated and unfiltered KL-expanded priors are summarized in Supplementary Table S1.

### Inference errors from KL-expanded priors are enriched among kinases with limited curated support

We next examined whether inference errors from KL-expanded priors were uniformly distributed across kinases. Directional error rates varied substantially among target kinases. Some kinases that were already evaluable using curated KSRs maintained relatively low error rates after KL expansion. In contrast, several kinases showed increasing error rates as more KL-derived relationships were incorporated. High-error kinase–library combinations were particularly frequent among newly evaluable kinases, especially in the broader KL95 and KL90 libraries (Fig. 2a).

**Figure 2.**
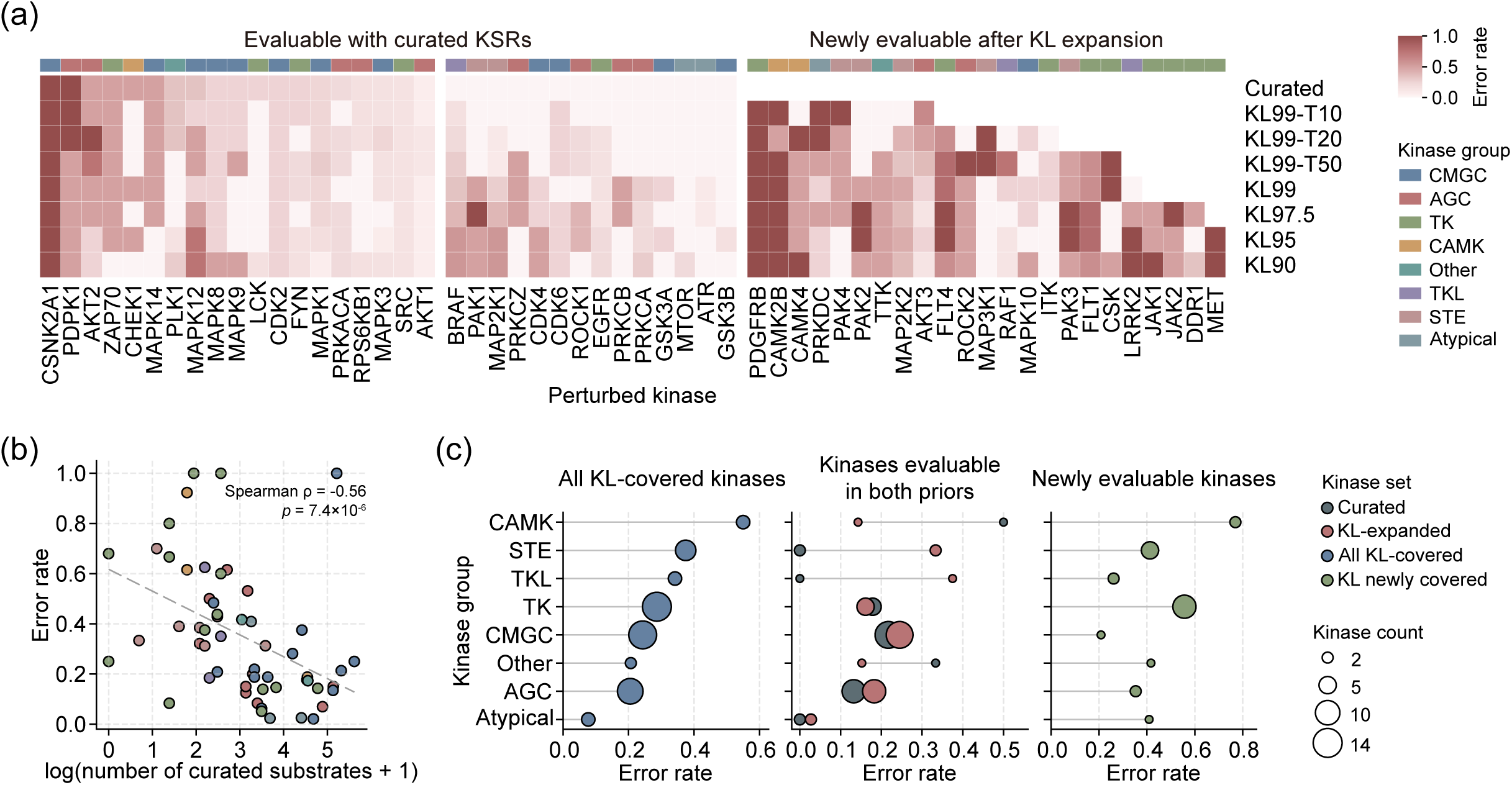
Inference errors from KL-expanded priors are enriched among kinases with limited curated support. **a,** Kinase-level directional error rates across KSR libraries. Columns correspond to evaluable kinases and rows correspond to KSR libraries. Kinases are grouped into those evaluable using the curated library and those newly evaluable after KL expansion. Kinase-group annotations are indicated above the heatmaps. Blank cells indicate kinases that were not evaluable under the corresponding library. **b,** Association between kinase-level directional error rate and the number of curated substrates for each kinase. Each point represents a kinase and is colored by kinase group. The x axis shows log-transformed curated substrate count plus one. Spearman *ρ* and p-value are shown. **c,** Directional error rates stratified by kinase group for all KL-covered kinases (left), kinases evaluable in both the curated and KL-expanded priors (middle), and kinases newly evaluable after KL expansion (right). In the middle panel, gray and red points indicate error rates from the curated and KL-expanded priors, respectively. Point size indicates the number of kinases in each group.

Directional error rates were inversely associated with curated substrate support. Across kinases covered by KL-expanded priors, the error rate decreased as the number of curated substrates increased (Spearman *ρ* = −0.56, *p* = 7.4×10^−6^; Fig. 2b). Kinases with strong curated support generally showed lower error rates, whereas several kinases with few or no curated substrates showed error rates above 0.6.

Error rates also differed across kinase groups. CAMK, STE, TKL and TK kinases showed higher average error rates than AGC and atypical kinases among KL-covered kinases (Fig. 2c, left). Among kinases evaluable in both curated and KL-expanded priors, KL expansion increased directional error rates for several kinase groups compared with the curated prior (Fig. 2c, middle). Newly evaluable kinases showed particularly high error rates in selected CAMK, STE, TKL and TK groups (Fig. 2c, right). These results indicate that the reliability of KL-expanded priors varies across the kinome. The largest errors were observed among kinases whose activity estimates depended primarily on KL-derived relationships, suggesting that additional cellular-context evidence may be required to improve their reliability.

### Protein-association constraints improve the accuracy–coverage balance of KL-expanded priors

We next investigated whether cellular-context information could refine KL-expanded priors and whether the benefit of filtering depended on specific types of network evidence. Curated KSRs were retained in all libraries, whereas KL-derived relationships were filtered using either predicted protein co-localization from DeepLoc[18] or protein-association constraints integrating STRING functional associations[19] with kinase-focused interactions from the Buljan et al. network[20]. For protein-association constraints, STRING400 and STRING700 retained KL-derived relationships supported by STRING combined scores ≥400 or ≥700, respectively, or by kinase–protein interactions from the Buljan et al. network (Fig. 3a; see Experimental Procedures). The resulting priors were evaluated using the same kinase activity inference framework (Supplementary Table S2).

**Figure 3.**
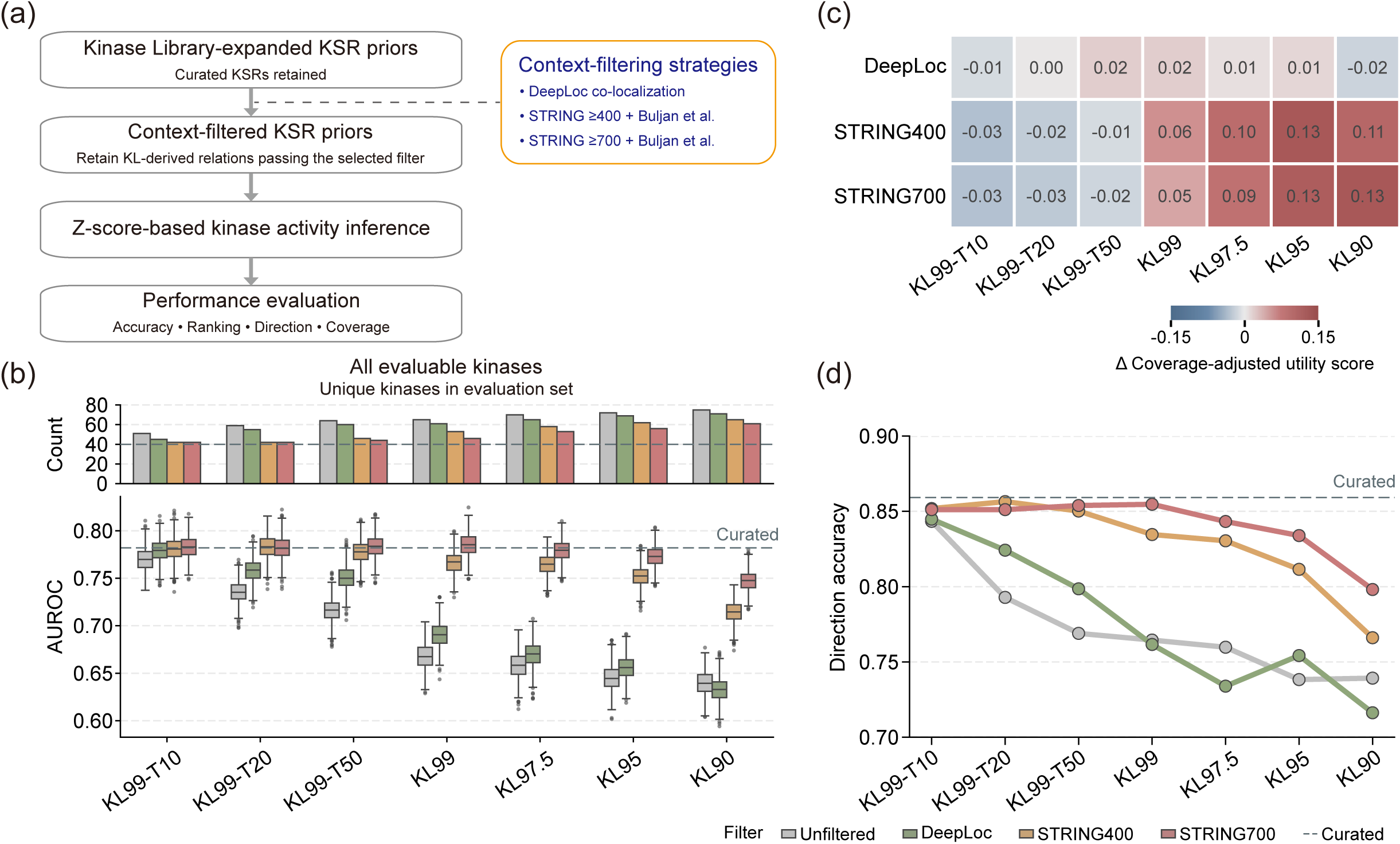
Cellular-context constraints improve the accuracy–coverage balance of KL-expanded priors. **a,** Overview of the cellular-context constraint strategy. Curated KSRs were retained unchanged, whereas KL-derived KSRs in each KL-expanded prior were constrained using DeepLoc-predicted protein co-localization or protein-association evidence. STRING400 and STRING700 retained KL-derived KSRs whose kinase–substrate pairs had STRING combined scores ≥400 or ≥700, respectively, or were supported by kinase-focused protein interactions reported by Buljan et al. The resulting context-constrained libraries were used for kinase activity inference using the weighted z-score formulation and evaluated by discrimination, rank-based recovery, directional accuracy and coverage. **b,** Evaluable kinases (top) and AUROC (bottom) across unfiltered and context-constrained KL-expanded priors. The dashed line indicates the median AUROC of the curated library. **c,** Heatmap showing the change in coverage-adjusted utility score after applying context constraints relative to the corresponding unfiltered KL-expanded prior at the same KL threshold. Positive values indicate an improved accuracy–coverage trade-off after filtering KL-derived KSRs. **d,** Directional accuracy across unfiltered and context-constrained KL-expanded priors. The dashed line indicates the directional accuracy of the curated library.

The effect of context filtering depended on the extent of KL expansion. Restricted libraries, particularly KL99-T10, showed limited changes after filtering, whereas broader KL-expanded priors showed substantial improvements after STRING-based filtering. For KL99, KL97.5, KL95 and KL90, both STRING400 and STRING700 improved AUROC relative to the corresponding unfiltered priors while maintaining substantially higher kinase coverage than the curated prior (Fig. 3b). In contrast, DeepLoc filtering produced smaller and less consistent improvements, indicating that protein-association evidence provided more effective constraints than broad subcellular co-localization for refining KL-derived KSRs. To determine whether the AUROC improvement was independent of differences in the evaluated kinase set, we compared all priors using only kinases with curated KSRs. The improvement after STRING-based filtering was largely retained under this controlled comparison (Supplementary Fig. S2a). Target-kinase ranking showed a similar pattern, with STRING filtering generally improving percentile ranks, particularly for the broader KL97.5, KL95 and KL90 libraries (Supplementary Fig. S2b–d).

While both STRING-based strategies improved inference performance, they differed in the balance between performance and kinase coverage. STRING700 generally achieved higher AUROC, whereas STRING400 retained more evaluable kinases. To quantify this trade-off, we calculated a coverage-adjusted utility score (see Experimental Procedures). DeepLoc produced limited changes in utility across most KL thresholds, whereas STRING400 and STRING700 increased utility for broader KL-expanded priors. The largest gains were observed for KL95 and KL90, with increases of approximately 0.11–0.13 relative to the corresponding unfiltered priors (Fig. 3c). STRING-based filtering also improved directional accuracy across broad KL-expanded priors (Fig. 3d), with improvements observed for both inhibitory and activating perturbations (Supplementary Fig. S2e). For KL90, directional accuracy increased from approximately 0.72 without filtering to approximately 0.77 and 0.80 after STRING400 and STRING700 filtering, respectively. These results demonstrate that protein-association constraints improve the accuracy–coverage balance of KL-derived KSRs.

To test whether similar gains could be obtained when network context was already incorporated into the prediction framework, we applied the same filtering strategies to the NetworKIN-expanded library[22]. Filtering produced only modest changes, with STRING700 showing a small improvement when evaluated on kinases with curated KSRs (Supplementary Fig. S3a,b; Supplementary Table S3). We next examined kinase-level responses to context filtering in KL-expanded priors. STRING400 and STRING700 reduced error rates for many kinase–library combinations, particularly in broad KL-expanded libraries, although some kinases showed little improvement or even increased error rates after filtering (Supplementary Fig. S3c–e). These results indicate that STRING-based filtering improves inference performance for broad KL-expanded priors while individual kinase responses remain heterogeneous.

Because STRING combined scores integrate multiple evidence channels, we further examined whether the benefit of STRING filtering was driven by literature-derived associations. Decomposition of STRING evidence showed that experimental and co-expression channels retained improvements comparable to the full STRING400 strategy, whereas text-mining and database-derived channels showed more variable effects (Supplementary Fig. S4a,b). STRING-only filtering recapitulated most of the overall coverage and performance gain of STRING400, whereas the Buljan network provided sparser but complementary support for selected kinases (Supplementary Fig. S4a–d). These results indicate that the benefit of STRING filtering is not primarily driven by literature-associated evidence. Among newly evaluable kinases, protein-association filters consistently improved inference performance compared with the unfiltered KL95 prior (Supplementary Fig. S4c,d). These analyses indicate that protein-association constraints improve KL-expanded priors by adding interaction-based evidence that is not solely dependent on literature-derived associations.

### Protein-association constraints reduce redundancy in KL-expanded kinase activity estimates

After establishing that protein-association constraints improve the accuracy–coverage balance of KL-expanded priors, we next asked whether expanded kinase coverage provided additional distinguishable activity information or primarily generated redundant estimates arising from overlapping KL-derived substrate assignments. Because kinases within the same family often share similar substrate-recognition patterns, we calculated pairwise Spearman *ρ* between kinase activity profiles across perturbation samples and compared the median correlations for within-family and between-family kinase pairs (see Experimental Procedures). Broad unfiltered KL-expanded priors, such as KL95, showed elevated within-family activity similarity while between-family correlations remained substantially lower, particularly for the more extensively expanded libraries. In contrast, STRING400 and STRING700 filtering attenuated the increase in within-family similarity and shifted the correlation patterns toward those observed with the curated prior (Fig. 4a). The same pattern was observed when the analysis was restricted to scoreable kinases with curated KSRs, indicating that the increased similarity was not explained solely by the addition of newly scoreable kinases (Supplementary Fig. S5a).

**Figure 4.**
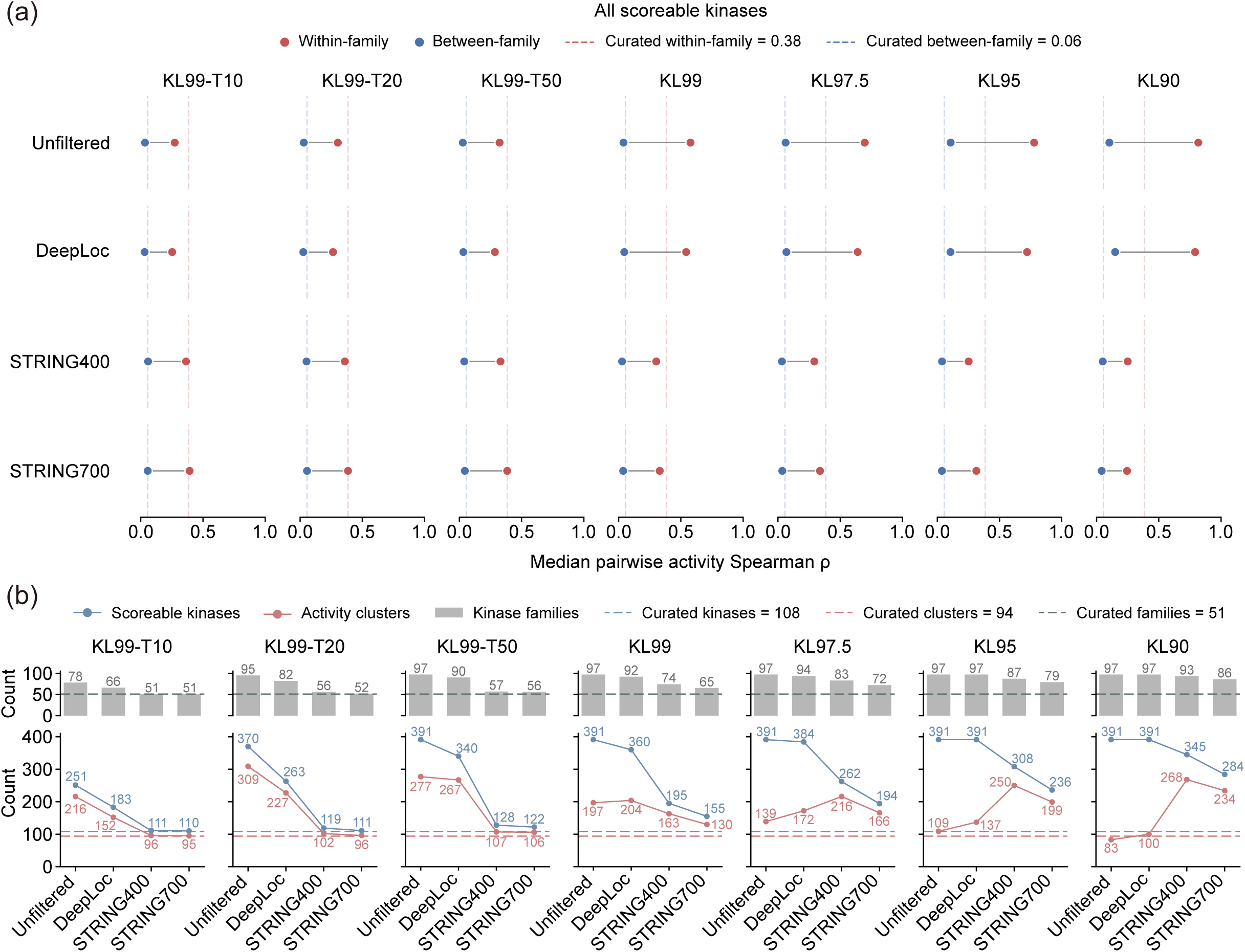
Context filtering reduces redundancy in KL-expanded kinase activity estimates. **a,** Median pairwise Spearman correlations (*ρ*) between kinase activity profiles for within-family and between-family kinase pairs across unfiltered and context-constrained KL-expanded priors. Dashed lines indicate the corresponding median correlations obtained using the curated KSR library. **b,** Kinase-family coverage (top) and numbers of scoreable kinases and correlation-defined activity clusters (bottom) across unfiltered and context-constrained KL-expanded priors. Dashed lines indicate the corresponding values obtained using the curated KSR library.

We next quantified the effective diversity of inferred kinase activity profiles using correlation-based activity clusters (see Experimental Procedures). Across KL-expanded priors, increases in the number of scoreable kinases were not accompanied by proportional increases in the number of distinguishable activity clusters (Fig. 4b). For example, although unfiltered KL95 expanded nominal coverage to 391 scoreable kinases, these kinases were grouped into only 109 activity clusters, indicating substantial redundancy among the expanded kinase activity estimates. In contrast, STRING400 filtering of KL95 retained 308 scoreable kinases distributed across 250 activity clusters while preserving broad kinase-family coverage (Fig. 4b; Supplementary Fig. S5b).

To investigate whether this redundancy was associated with overlap in the underlying substrate sets, we compared pairwise KSR overlap with kinase activity similarity (see Experimental Procedures). As KL expansion became broader, unfiltered priors showed an increasingly strong association between phosphosite-level KSR Jaccard similarity and activity-profile similarity, consistent with overlapping motif-derived substrate assignments contributing to correlated activity estimates. This association remained elevated after DeepLoc filtering but was substantially reduced by STRING400 and STRING700, approaching the level observed for the curated prior (Supplementary Fig. S5c). To facilitate kinase-level interpretation, we additionally compiled per-kinase reliability annotations integrating complementary evidence on substrate support, activity distinguishability, and benchmark-derived reliability across KSR priors (Supplementary Table S4). These analyses indicate that protein-association filtering reduces the redundancy introduced by broad KL expansion while preserving substantial kinase coverage.

### Protein-association-supported substrates show stronger agreement with expected perturbation direction

We next examined whether the improvement produced by STRING filtering was evident at the substrate level. KL-derived phosphosites quantified in each perturbation experiment were classified as STRING-supported or STRING-unsupported, and their phosphorylation changes were compared across representative kinase inhibition and activation experiments spanning different degrees of KL expansion.

In HL60 cells treated with the PDPK1 inhibitor GSK2334470, PDPK1 activity was expected to decrease. The KL99 prior assigned 99 quantified substrate sites to PDPK1, of which 13 were supported by STRING400 and 86 were unsupported. STRING400-supported substrates showed a stronger shift toward decreased phosphorylation than the unsupported substrates (Fig. 5a). The STRING400-supported network included GSK3B S9, PLK1 T210 and RPS6KA-family sites, many of which showed phosphorylation changes consistent with PDPK1 inhibition (Fig. 5b).

**Figure 5.**
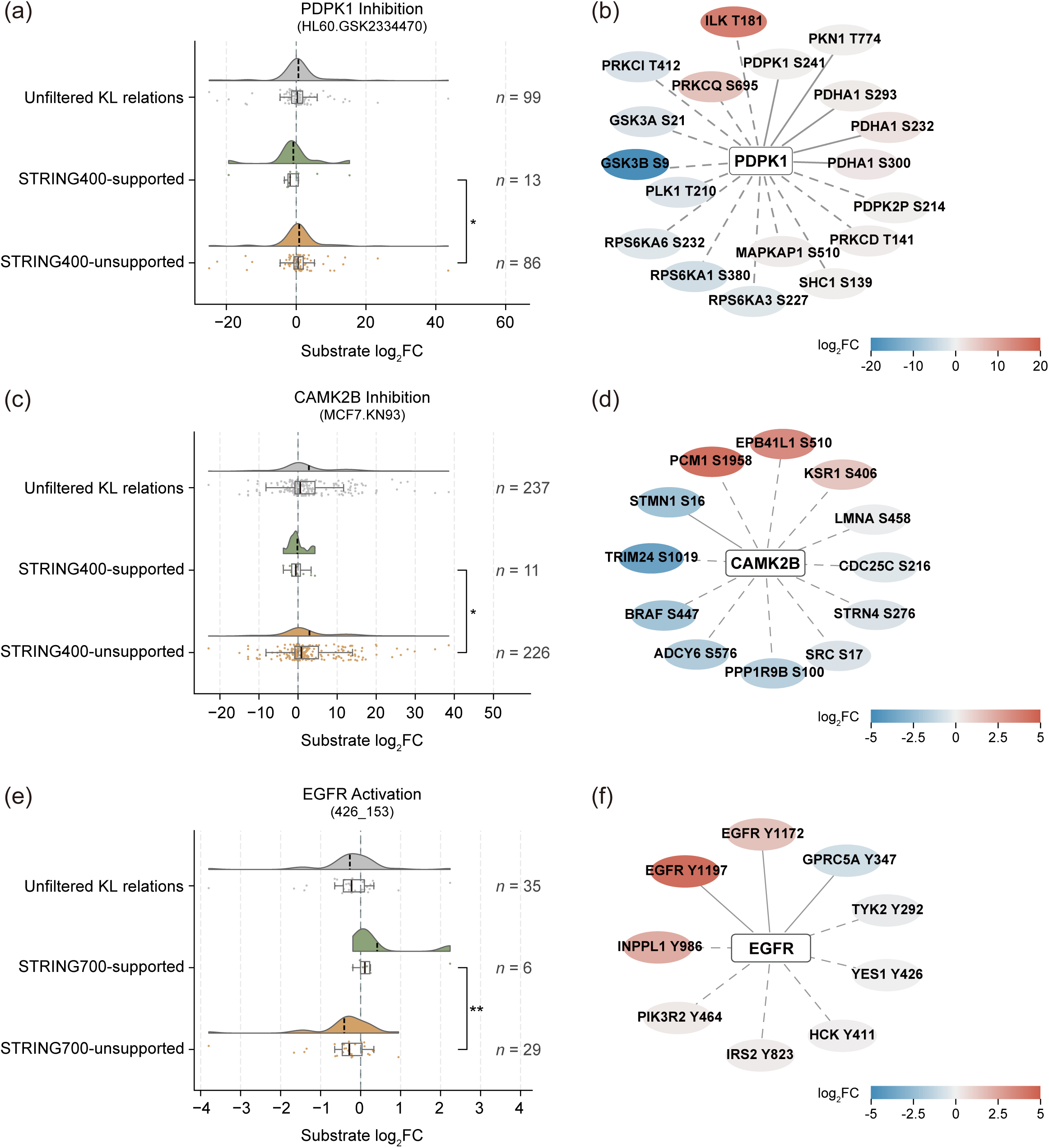
Protein-association-supported KL substrate sites show stronger agreement with perturbation-induced phosphorylation changes. **a,** Phosphorylation changes (log2 fold changes) of KL-derived PDPK1 substrate sites in HL60 cells treated with the PDPK1 inhibitor GSK2334470. Sites from the unfiltered KL99 library are shown together with STRING400-supported and STRING400-unsupported subsets. The number of sites in each group is indicated. **b,** Network view of PDPK1-linked curated substrate sites (solid edges) and KL-derived substrate sites retained in the STRING400-constrained KL99 library (dashed edges) for the same perturbation sample shown in **a**. **c,** Phosphorylation changes (log2 fold changes) of KL-derived CAMK2B substrate sites in MCF7 cells treated with the CAMK2B inhibitor KN93. Sites from the unfiltered KL97.5 library are shown together with STRING400-supported and STRING400-unsupported subsets. The number of sites in each group is indicated. **d,** Network view of CAMK2B-linked curated substrate sites (solid edges) and KL-derived substrate sites retained in the STRING400-constrained KL97.5 library (dashed edges) for the same perturbation sample shown in **c**. **e,** Phosphorylation changes (log2 fold changes) of KL-derived EGFR substrate sites in EGF-stimulated HeLa cells. Sites from the unfiltered KL90 library are shown together with STRING700-supported and STRING700-unsupported subsets. The number of sites in each group is indicated. **f,** Network view of EGFR-linked curated substrate sites (solid edges) and KL-derived substrate sites retained in the STRING700-constrained KL90 library (dashed edges) for the same perturbation sample shown in **e**. Substrate-site nodes are colored according to phosphosite log2 fold change. Networks in **b**, **d** and **f** were visualized in Cytoscape. Statistical comparisons between STRING-supported and STRING-unsupported groups were performed using one-sided Mann– Whitney U tests in the direction expected from the annotated kinase perturbation. \**p* < 0.05, \*\**p* < 0.01.

A similar pattern was observed for CAMK2B inhibition in KN93-treated MCF7 cells. The KL97.5 prior assigned 237 quantified substrate sites to CAMK2B, including 11 STRING400-supported and 226 unsupported sites. The supported subset showed a stronger shift toward decreased phosphorylation than the unsupported subset (Fig. 5c). The corresponding network included STMN1, BRAF, ADCY6 and PPP1R9B (Fig. 5d).

We also examined EGFR activation after EGF stimulation in HeLa cells. The KL90 prior assigned 35 quantified substrate sites to EGFR, of which 6 were supported by STRING700 and 29 were unsupported. STRING700-supported substrates showed a stronger shift toward increased phosphorylation than the unsupported subset (Fig. 5e). The retained network included EGFR autophosphorylation sites and signaling proteins including INPPL1, PIK3R2 and IRS2 (Fig. 5f). Across these examples, protein-association filtering preferentially retained motif-compatible substrates whose phosphorylation changes were more consistent with the expected direction of kinase perturbation.

### Accuracy–coverage landscape defines operating regimes for KL-expanded priors

We summarized all threshold–filter combinations using an accuracy–coverage map. The curated prior supported activity estimates for 108 kinases and achieved an AUROC of approximately 0.78, whereas KL99-T10 maintained similar accuracy but provided only limited additional kinase coverage. In contrast, broadly expanded unfiltered and DeepLoc-filtered libraries covered approximately 350–400 kinases but generally showed AUROC values below 0.70 (Fig. 6a).

**Figure 6.**
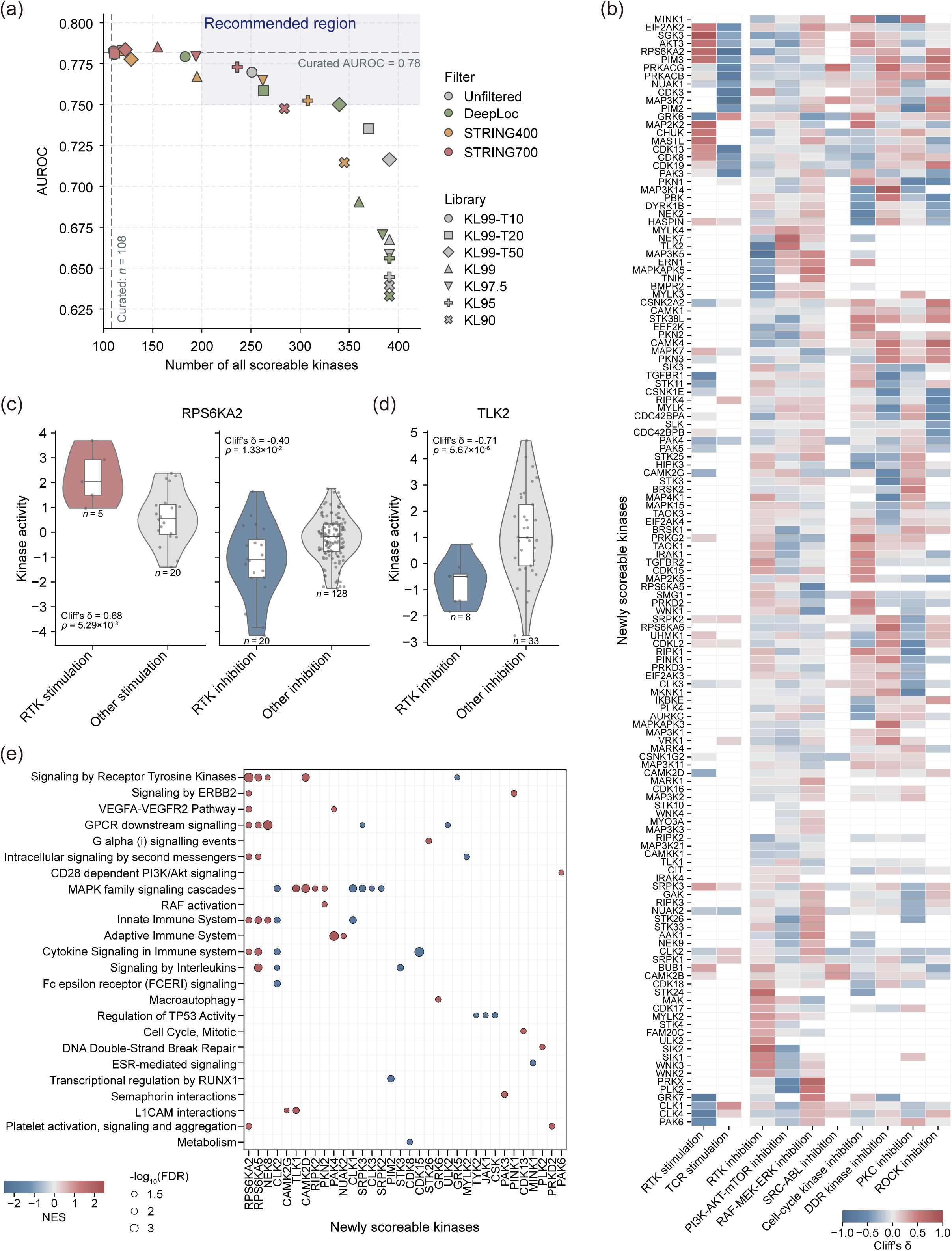
Accuracy–coverage landscape identifies KL95–STRING400 as a practical prior for biological interpretation. **a,** Accuracy is represented by median AUROC, and coverage by the number of scoreable kinases. Each point represents a library–filter combination. The shaded region indicates the recommended operating region with at least 200 scoreable kinases and AUROC ≥0.75. The horizontal dashed line indicates the median AUROC of the curated library, and the vertical dashed line indicates the number of scoreable kinases using the curated library. **b,** Heatmap showing perturbation-class activity patterns of newly scoreable kinases. For each kinase and focal perturbation class, inferred activities were compared with those in all other samples with the same perturbation direction. Colors indicate Cliff’s δ, with positive and negative values representing higher and lower activity in the focal perturbation class, respectively. Blank cells indicate comparisons that did not meet the minimum sample requirements. **c,** Inferred RPS6KA2 activity following RTK stimulation compared with other stimulation conditions (left) and following RTK inhibition compared with other inhibition conditions (right). Sample sizes are shown below each group. Statistical comparisons were performed using two-sided Brunner–Munzel tests, with Cliff’s δ shown as the effect size. **d,** Inferred TLK2 activity following RTK inhibition compared with other inhibition conditions. Sample sizes are shown below each group. Statistical comparisons were performed using a two-sided Brunner–Munzel test, with Cliff’s δ shown as the effect size. **e,** Representative Reactome pathways significantly associated with newly scoreable kinases by preranked gene-set enrichment analysis (FDR *q* < 0.05). Colors indicate normalized enrichment scores (NES), and point size indicates −log_10_(FDR *q*).

STRING-filtered libraries occupied an intermediate region, retaining substantially broader kinase coverage while showing higher inference accuracy. Several STRING400- and STRING700-filtered priors achieved AUROC values above 0.75 while covering more than 200 kinases. STRING400 generally preserved more kinase coverage, whereas STRING700 provided higher accuracy under more stringent filtering. We therefore defined a practical operating region as priors with at least 200 scoreable kinases and AUROC ≥0.75 (Fig. 6a). Within this region, KL95-STRING400 provided broad kinase coverage while maintaining high inference accuracy and was selected for subsequent biological analyses.

### KL95-STRING400 enables biological interpretation of newly scoreable kinases

To determine whether the expanded kinase coverage provided biologically interpretable activity estimates, we focused on the 200 kinases that were scoreable using KL95-STRING400 but not using the curated KSR library. Among these newly scoreable kinases, 59 were classified as dark kinases by the Illuminating the Druggable Genome (IDG) program[24], compared with 7 of the 108 kinases scoreable using curated KSRs (Fisher’s exact test, *p* = 9.3×10^−7^; Supplementary Fig. S6a; see Experimental Procedures). Thus, the additional coverage provided by KL95-STRING400 was significantly enriched for understudied kinases.

We next examined whether these newly scoreable kinases showed systematic activity differences across perturbation contexts. The 222 perturbation experiments were grouped into major perturbation classes, including RTK stimulation, TCR stimulation, and several kinase-inhibition categories, with sufficiently represented classes retained for subsequent analysis (Supplementary Fig. S6b; Supplementary Table S5; see Experimental Procedures). Newly scoreable kinases displayed distinct activity patterns across these perturbation classes (Fig. 6b), indicating condition-associated variation that could not be assessed using curated KSRs alone. For example, RPS6KA2 showed increased inferred activity following RTK stimulation and decreased activity under RTK inhibition (Fig. 6c), consistent with previous evidence that EGF/EGFR signaling activates RPS6KA2 through the MEK–ERK cascade[29]. TLK2, an IDG-defined dark kinase, also exhibited reduced activity following RTK inhibition compared with other inhibition conditions (Fig. 6d). Previous studies have linked TLK2 to EGFR/HER2/SRC/FAK signaling[30], supporting an association between TLK2 activity and RTK-related signaling.

To further characterize the functional context of newly scoreable kinases, we correlated their activity profiles with those of kinases scoreable using curated KSRs and used the resulting rankings for preranked gene-set enrichment analysis (see Experimental Procedures). Same-family kinases represented only a small fraction of the curated-scoreable reference kinases, with a median proportion of 1.3% across newly scoreable kinases (Supplementary Fig. S6c). Several kinases showed significant associations with established signaling programs (Fig. 6e; Supplementary Table S6). For example, RPS6KA2 showed a strong positive association with RTK signaling (NES = 2.03, FDR *q* < 0.001), consistent with its activity pattern observed in the perturbation-class analysis. Additionally, the dark kinase PAK6 was positively associated with CD28-dependent PI3K/Akt signaling (NES = 1.66, FDR *q* = 0.04), consistent with previous evidence that AKT inhibition reduces PAK6 phosphorylation[31]. These analyses show that KL95-STRING400 extends kinase activity inference to kinases inaccessible using curated KSRs alone, including a substantial fraction of the understudied kinome, while yielding biologically interpretable activity patterns.

## Discussion

Our benchmark defines the limitations and opportunities of KL-based KSR expansion. The additional relationships increase kinase coverage, but they can also reduce the regulatory specificity of kinase substrate sets. This reflects the different evidence encoded by curated and motif-derived KSRs. Curated relationships provide prior evidence of intracellular regulation, whereas KL predictions quantify intrinsic sequence compatibility[12, 13]. These evidence types are complementary, but not interchangeable.

The higher error rates among kinases with limited curated substrate support are particularly relevant to less characterized kinases within the human kinome[10, 11]. For well-studied kinases, curated substrates can anchor the inferred activity signal. For poorly characterized kinases, the estimate may depend largely on predicted substrates. Our results further show that increased kinase coverage does not necessarily provide proportionally more distinguishable activity information. Broad KL expansion increased within-family activity similarity and produced redundant activity profiles. Protein-association filtering reduced this redundancy and retained more distinguishable activity patterns. Newly gained scoreability should therefore be distinguished from both reliable inference and increased information content.

Protein-association evidence provided a more informative constraint than predicted co-localization. Broad compartment annotations can exclude clearly incompatible pairs but offer limited specificity within large cellular compartments[14]. They also do not capture dynamic translocation or subcompartment organization[14]. Protein-association evidence places candidate kinase–substrate pairs in a more specific functional context. Importantly, the improvement was also observed when STRING evidence was restricted to experimental and co-expression channels, indicating that the benefit was not primarily driven by literature-derived associations. At the substrate level, STRING-supported relationships also showed phosphorylation changes that were more consistent with the expected perturbation direction. STRING support, however, does not prove direct phosphorylation of a particular site. The retained relationships should remain classified as prioritized candidates.

The comparison between STRING400 and STRING700 further illustrates that prior selection depends on the analytical objective. Stringent filtering may be preferable when inference reliability is prioritized, whereas moderate filtering may be more useful when broader kinome coverage is required. The accuracy–coverage trade-off therefore provides a practical framework for selecting KL-expanded priors for different applications.

An important implication of context-filtered KL expansion is its potential to extend kinase activity inference into less characterized regions of the kinome, where curated substrate evidence is sparse. The enrichment of dark kinases among newly scoreable kinases suggests that this gain in coverage preferentially reaches kinases that are difficult to evaluate using curated priors alone. The interpretable perturbation and pathway associations observed for kinases such as RPS6KA2, TLK2 and PAK6 further support the use of this strategy for hypothesis generation. These associations, however, should not be considered direct validation of individual kinase activities or KSRs.

Several limitations remain. The perturbation benchmark covers only part of the human kinome and is biased toward kinases with available phosphoproteomic perturbation data. Inhibitor-based perturbations may also involve off-target or indirect pathway effects[6, 7]. Moreover, the contextual evidence used here is generic rather than sample-specific and does not explicitly capture protein abundance, cell type, dynamic localization or condition-specific interactions. Regulatory phosphorylation of kinases may represent an additional layer of kinase activity control[32]. Incorporating such information, together with kinase-specific constraints on substrate recognition, may further improve inference. Overall, protein-association evidence provides a practical way to refine motif-derived KSR priors while preserving broad kinase coverage.

## Data Availability

All public datasets used in this study are available from their original sources. The benchmark phosphoproteomic matrix, perturbation annotations, and curated KSR set used in this study are publicly available through the benchmarKIN R package (https://github.com/saezlab/benchmarKIN). NetworKIN predictions were obtained from the associated benchmarKIN Zenodo record (https://zenodo.org/records/14824013). Kinase Library predictions were generated using the Kinase Library Python package (https://github.com/TheKinaseLibrary/kinase-library). DeepLoc subcellular localization predictions were performed using the DeepLoc 2.1 web server (https://services.healthtech.dtu.dk/services/DeepLoc-2.1/). Protein-association data were obtained from STRING version 12.0 (https://stringdb-downloads.org/download/protein.links.detailed.v12.0/9606.protein.links.detailed.v12.0.txt.gz).

Processed data generated in this study are available at https://github.com/ZhangMenghuan-Tongji/KLAIC, including KL-expanded KSR priors, the NetworKIN-expanded prior, context-filtered priors, inferred kinase activity matrices, and evaluation summary tables.

## Code availability

All source code and custom scripts used to construct KL-expanded and context-constrained libraries, perform z-score kinase activity inference, calculate evaluation metrics, and reproduce the figures are available at https://github.com/ZhangMenghuan-Tongji/KLAIC.

## Acknowledgments

This work was supported by the National Natural Science Foundation of China (32670906 to M.Z.).

## Author Contributions

M.Z. conceived and supervised the study, designed the analytical framework, interpreted the results and wrote the manuscript. Y.Z. performed data processing, benchmark construction, kinase activity inference, context filtering and evaluation analyses, developed the software package and prepared the figures. Both authors reviewed and approved the final manuscript.

