## Supplementary Information for "Improving phosphoproteomic kinase activity inference by contextual refinement of Kinase Library relationships"

### Supplementary Tables

Supplementary Table S1. Benchmark summary for curated and unfiltered KL-expanded libraries.

Supplementary Table S2. Benchmark summary for context-constrained KL-expanded libraries.

Supplementary Table S3. Benchmark summary for unfiltered and context-constrained NetworKIN-expanded libraries.

Supplementary Table S4. Per-kinase activity-cluster and reliability annotations across KSR libraries.

Supplementary Table S5. Perturbation-class analysis of newly scoreable kinases.

Supplementary Table S6. Reactome GSEA results for newly scoreable kinases.

### Supplementary Figures

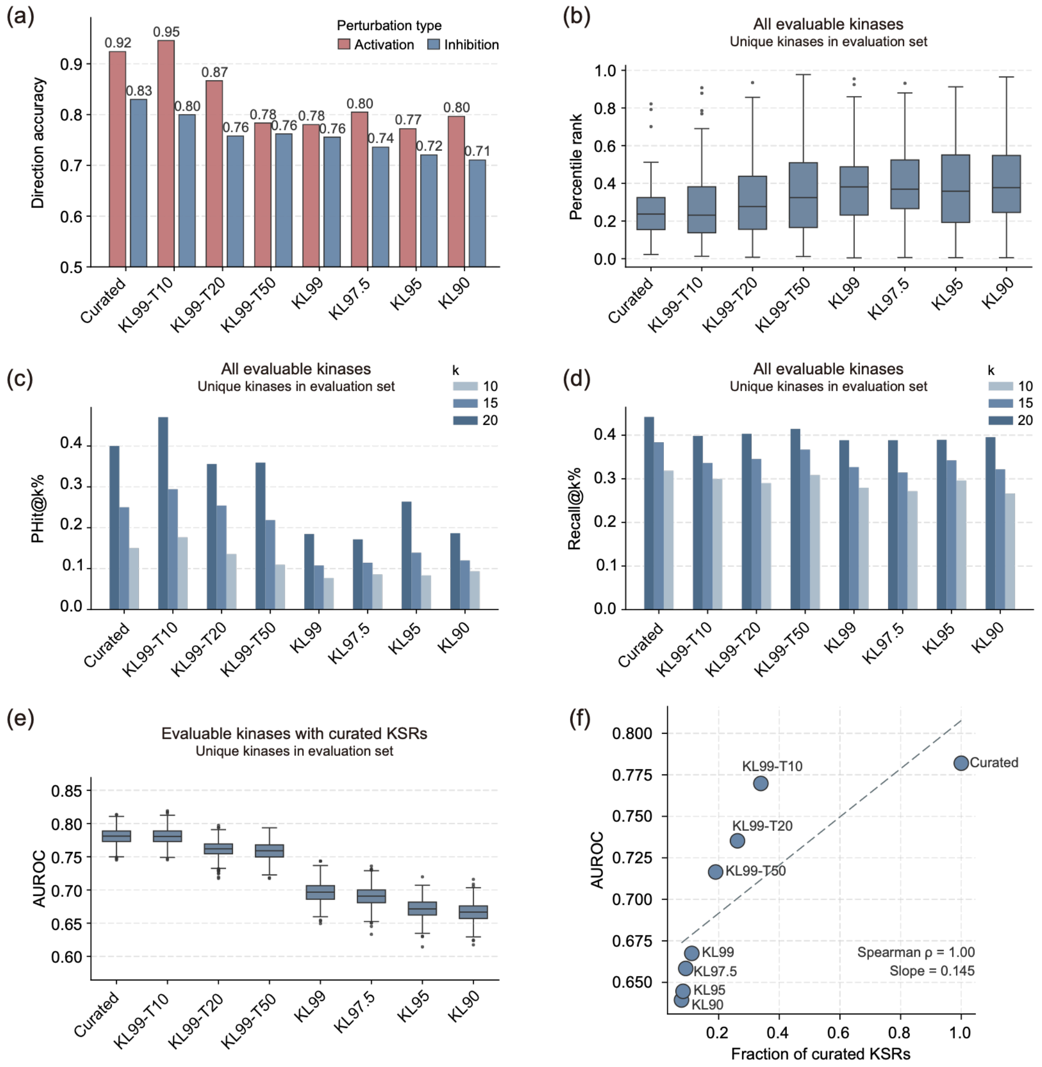


**Supplementary Figure S1. Additional evaluation of kinase activity inference performance using KL-expanded priors.**

**a,** Directional accuracy stratified by perturbation type across the curated and Kinase Library (KL)-expanded priors. Inhibition and activation events were evaluated separately.

**b,** Percentile ranks of evaluable target kinases among all kinases with inferred activity. Lower percentile ranks indicate better target-kinase recovery.

**c,** PHit@k% across kinase–substrate relationship (KSR) libraries, defined as the fraction of evaluable target kinases whose mean percentile rank across samples fell within the top k%, with k = 10, 15 or 20.

**d,** Recall@k% across KSR libraries, defined as the mean fraction of evaluable target kinases recovered within the top k% of kinases ranked by inferred activity in each sample, with k = 10, 15 or 20.

**e,** Area under the receiver operating characteristic curve (AUROC) when evaluation was restricted to kinases evaluable using the curated KSR library.

**f,** Association between AUROC and the fraction of KSRs in each library that were curated relationships. Spearman $\rho$ and the fitted slope are shown.

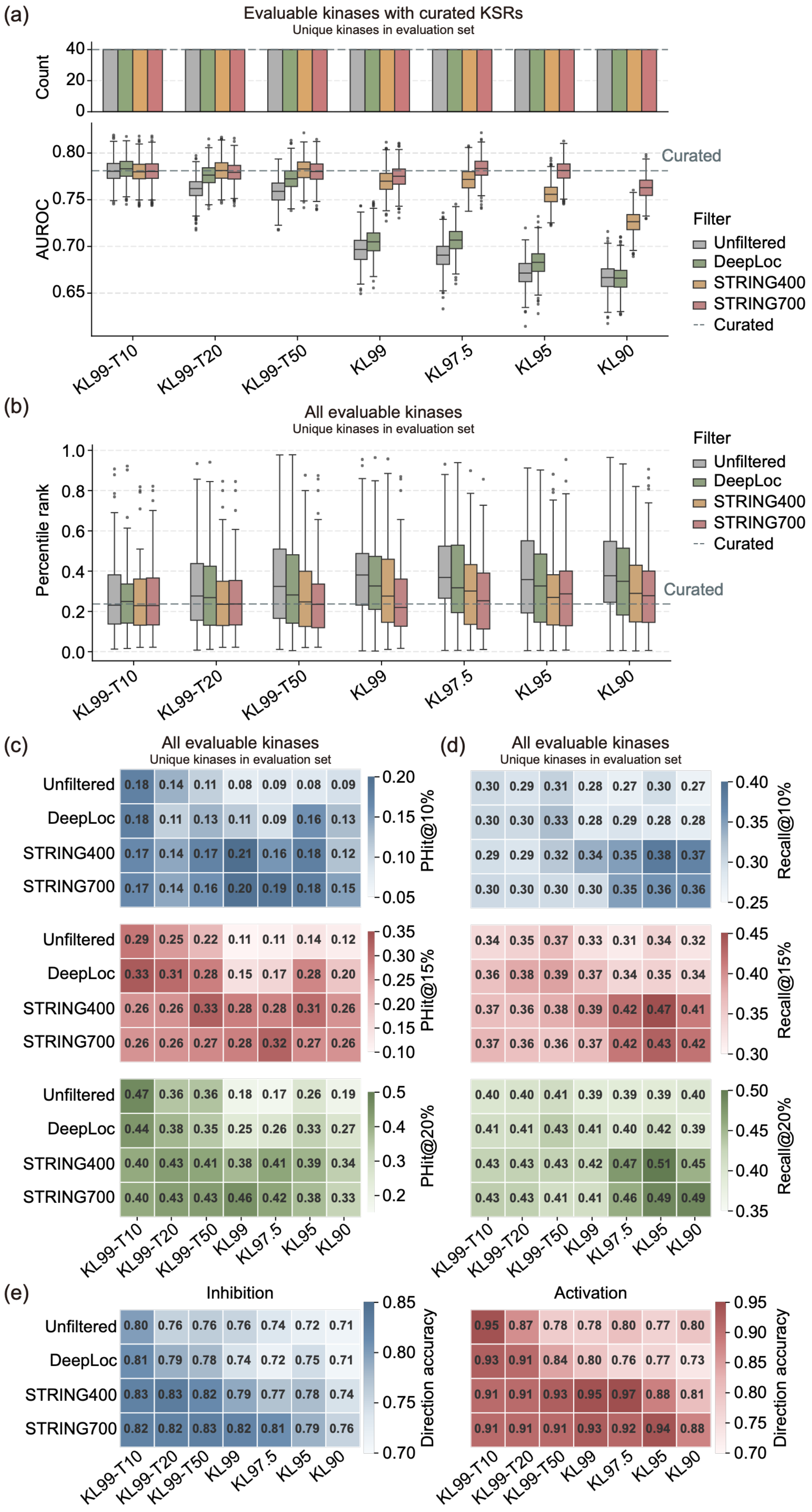

(a)
Evaluable kinases with curated KSRs
Unique kinases in evaluation set
Count
40
20
0
AUROC
0.80
0.75
0.70
0.65
Curated
Filter
Unfiltered
DeepLoc
STRING400
STRING700
Curated
KL99-T10
KL99-T20
KL99-T50
KL99
KL97.5
KL95
KL90
(b)
All evaluable kinases
Unique kinases in evaluation set
Percentile rank
1.0
0.8
0.6
0.4
0.2
0.0
Curated
Filter
Unfiltered
DeepLoc
STRING400
STRING700
Curated
KL99-T10
KL99-T20
KL99-T50
KL99
KL97.5
KL95
KL90
(c)
All evaluable kinases
Unique kinases in evaluation set
Unfiltered 0.18 0.14 0.11 0.08 0.09 0.08 0.09
DeepLoc 0.18 0.11 0.13 0.11 0.09 0.16 0.13
STRING400 0.17 0.14 0.17 0.21 0.16 0.18 0.12
STRING700 0.17 0.14 0.16 0.20 0.19 0.18 0.15
PHit@10%
0.20
0.15
0.10
0.05
Unfiltered 0.29 0.25 0.22 0.11 0.11 0.14 0.12
DeepLoc 0.33 0.31 0.28 0.15 0.17 0.28 0.20
STRING400 0.26 0.26 0.33 0.28 0.28 0.31 0.26
STRING700 0.26 0.26 0.27 0.28 0.32 0.27 0.26
PHit@15%
0.35
0.30
0.25
0.20
0.15
0.10
Unfiltered 0.47 0.36 0.36 0.18 0.17 0.26 0.19
DeepLoc 0.44 0.38 0.35 0.25 0.26 0.33 0.27
STRING400 0.40 0.43 0.41 0.38 0.41 0.39 0.34
STRING700 0.40 0.43 0.43 0.46 0.42 0.38 0.33
PHit@20%
0.5
0.4
0.3
0.2
KL99-T10
KL99-T20
KL99-T50
KL99
KL97.5
KL95
KL90
(d)
All evaluable kinases
Unique kinases in evaluation set
0.30 0.29 0.31 0.28 0.27 0.30 0.27
0.30 0.30 0.33 0.28 0.29 0.28 0.28
0.29 0.29 0.32 0.34 0.35 0.38 0.37
0.30 0.30 0.30 0.30 0.35 0.36 0.36
Recall@10%
0.40
0.35
0.30
0.25
0.34 0.35 0.37 0.33 0.31 0.34 0.32
0.36 0.38 0.39 0.37 0.34 0.35 0.34
0.37 0.36 0.38 0.39 0.42 0.47 0.41
0.37 0.36 0.36 0.37 0.42 0.43 0.42
Recall@15%
0.45
0.40
0.35
0.30
0.40 0.40 0.41 0.39 0.39 0.39 0.40
0.41 0.41 0.43 0.41 0.40 0.42 0.39
0.43 0.43 0.43 0.42 0.47 0.51 0.45
0.43 0.43 0.41 0.41 0.46 0.49 0.49
Recall@20%
0.50
0.45
0.40
0.35
KL99-T10
KL99-T20
KL99-T50
KL99
KL97.5
KL95
KL90
(e)
Inhibition
Unfiltered 0.80 0.76 0.76 0.76 0.74 0.72 0.71
DeepLoc 0.81 0.79 0.78 0.74 0.72 0.75 0.71
STRING400 0.83 0.83 0.82 0.79 0.77 0.78 0.74
STRING700 0.82 0.82 0.83 0.82 0.81 0.79 0.76
Direction accuracy
0.85
0.80
0.75
0.70
KL99-T10
KL99-T20
KL99-T50
KL99
KL97.5
KL95
KL90
Activation
0.95 0.87 0.78 0.78 0.80 0.77 0.80
0.93 0.91 0.84 0.80 0.76 0.77 0.73
0.91 0.91 0.93 0.95 0.97 0.88 0.81
0.91 0.91 0.91 0.93 0.92 0.94 0.88
Direction accuracy
0.95
0.90
0.85
0.80
0.75
0.70
KL99-T10
KL99-T20
KL99-T50
KL99
KL97.5
KL95
KL90

**Supplementary Figure S2. Additional evaluation of context-filtered KL priors.**

**a,** Number of evaluable kinases (top) and AUROC (bottom) when evaluation was restricted to kinases evaluable using the curated KSR library. The dashed line indicates the median AUROC of the curated library.

**b,** Percentile ranks of evaluable target kinases across unfiltered and context-constrained KL-expanded priors. Lower percentile ranks indicate better target-kinase recovery. The dashed line indicates the median percentile rank of the curated library.

**c,** PHit@k% across unfiltered and context-constrained KL-expanded priors, with k = 10, 15 or 20.

**d,** Recall@k% across unfiltered and context-constrained KL-expanded priors, with k = 10, 15 or 20.

**e,** Directional accuracy stratified by perturbation type across unfiltered and context-constrained KL-expanded priors. Inhibition (left) and activation (right) events were evaluated separately.

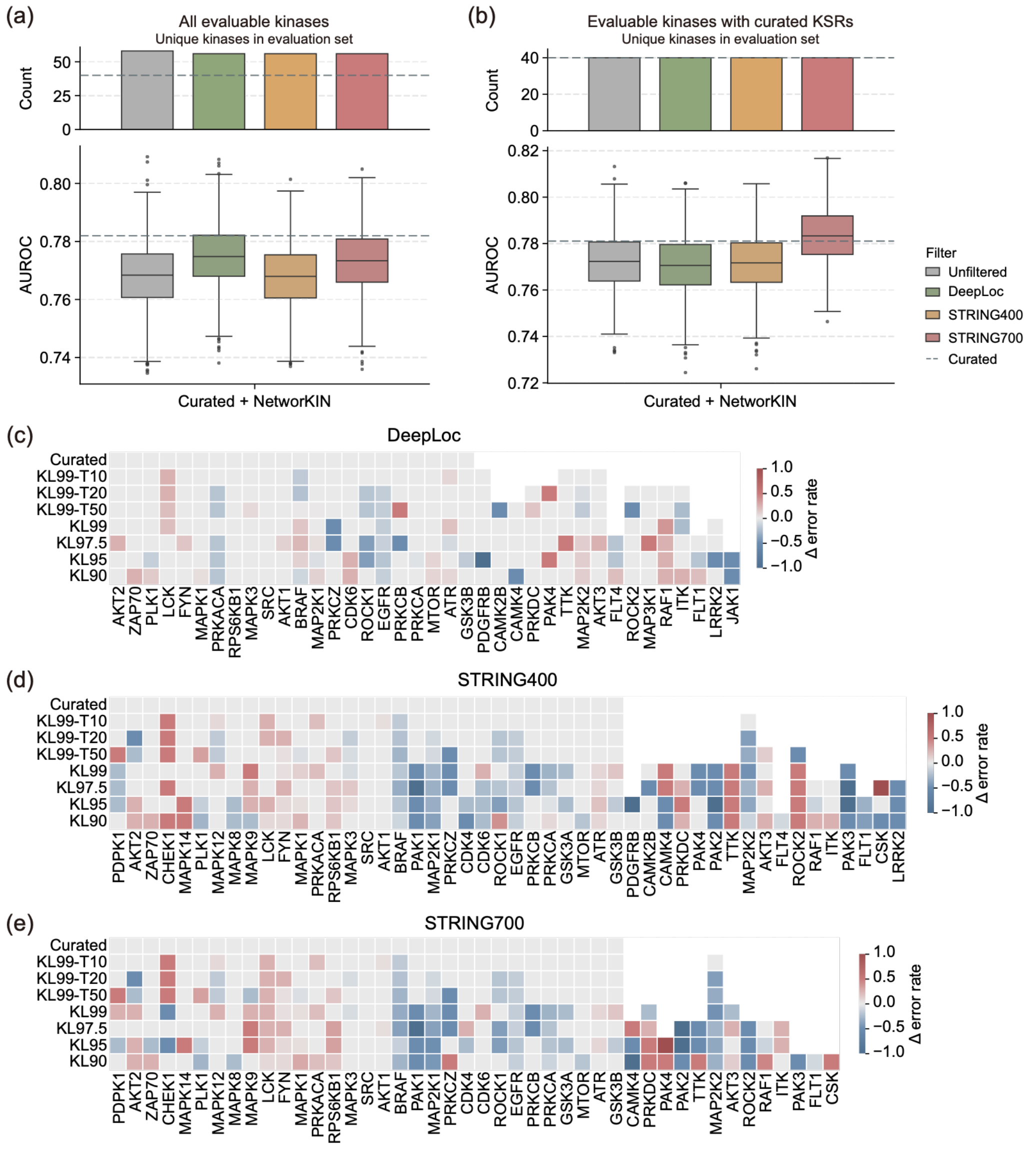


**Supplementary Figure S3. Effect of context filtering on NetworKIN-expanded library and kinase-level inference errors.**

**a–b,** Number of evaluable kinases (top) and AUROC (bottom) for the NetworKIN-expanded library using all evaluable kinases **(a)** and kinases evaluable with the curated library **(b)**. NetworKIN-derived KSRs were either left unfiltered or subjected to the same DeepLoc, STRING400 or STRING700 constraint strategies. The dashed line indicates the median AUROC of the curated library.

**c–e,** Changes in kinase-level directional error rates after applying DeepLoc **(c)**, STRING400 **(d)** or STRING700 **(e)** constraints to KL-expanded priors. Rows correspond to different KL-expanded libraries, and columns correspond to evaluable target kinases. Δ error rate was calculated relative to the corresponding unfiltered library at the same KL threshold. Negative values indicate reduced directional error after context filtering, whereas positive values indicate increased error. Blank cells indicate kinase–library combinations for which no directional error rate could be calculated.

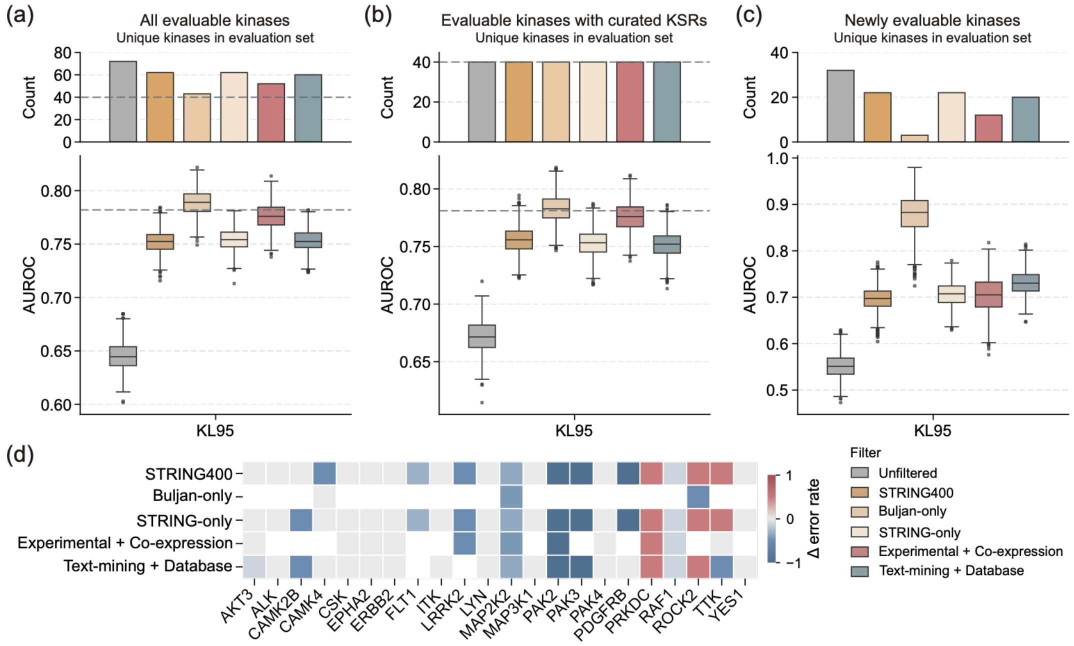


**Supplementary Figure S4. STRING evidence decomposition reveals complementary contributions to kinase activity inference.**

**a,** Number of evaluable kinases (top) and AUROC (bottom) for the unfiltered KL95 prior and KL95 priors filtered using STRING400, Buljan-only, STRING-only, the experimental and co-expression subset, or the text-mining and curated-database subset. STRING400 retained KL-derived KSRs supported by either a STRING combined score ≥400 or the Buljan et al. interaction network. Dashed lines indicate the corresponding values obtained using the curated KSR library.

**b,** Number of evaluable kinases (top) and AUROC (bottom) for the same filtering strategies when evaluation was restricted to target kinases evaluable using curated KSRs. Dashed lines indicate the corresponding values obtained using the curated KSR library.

**c,** Number of evaluable kinases (top) and AUROC (bottom) for newly evaluable kinases, defined as target kinases that were not evaluable using the curated KSR library but became evaluable after KL95 expansion.

**d,** Changes in kinase-level directional error rates after filtering relative to the unfiltered KL95 prior. Columns correspond to perturbed target kinases and rows to filtering strategies. Negative values indicate reduced directional error after filtering, whereas positive values indicate increased error. Blank cells indicate kinase–filter combinations for which a directional error rate could not be calculated.

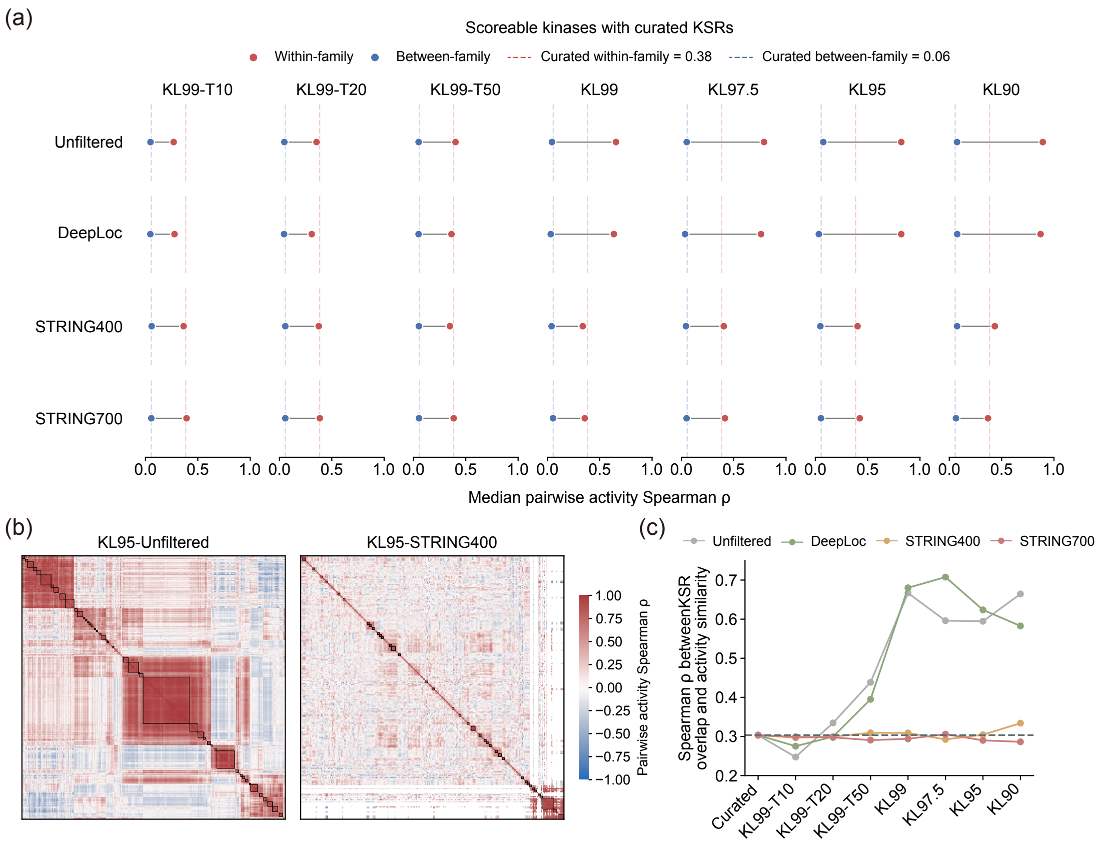


**Supplementary Figure S5. Additional analyses of activity redundancy across KL-expanded and context-filtered priors.**

**a,** Median pairwise Spearman correlations ($\rho$) between kinase activity profiles for within-family and between-family kinase pairs across unfiltered and context-constrained KL-expanded priors, restricted to kinases that were also scoreable using the curated KSR library. Dashed lines indicate the corresponding median correlations obtained using the curated KSR library.

**b,** Pairwise kinase activity correlation matrices for the KL95-unfiltered and KL95-STRING400 priors. Kinases are ordered by complete-linkage hierarchical clustering using $1-\rho$ as the correlation distance. Black outlines indicate correlation-defined activity clusters corresponding to a Spearman $\rho$ threshold of 0.7.

**c,** Association between KSR overlap and kinase activity similarity across unfiltered and context-constrained KL-expanded priors. KSR overlap was quantified using phosphosite-level Jaccard similarity, and activity similarity was quantified using pairwise Spearman $\rho$ between kinase activity profiles. The dashed line indicates the corresponding value obtained using the curated KSR library.

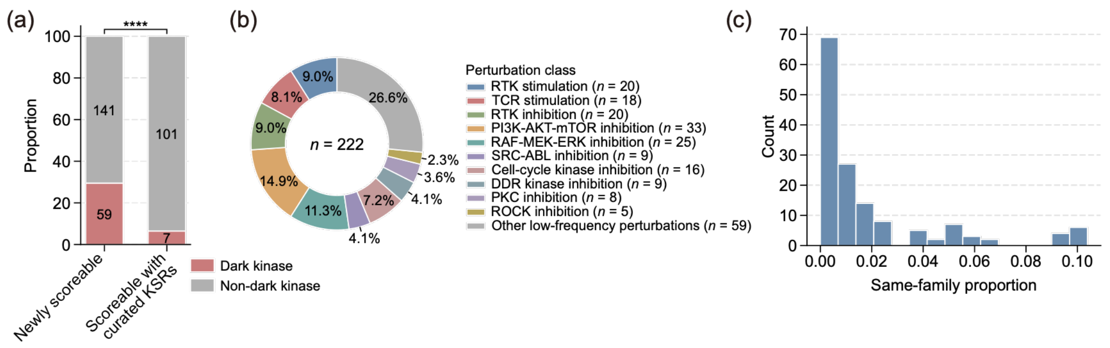


**Supplementary Figure S6. Characterization of newly scoreable kinases and perturbation contexts.**

**a,** Proportions of dark and non-dark kinases among newly scoreable kinases and kinases scoreable using curated KSRs. Dark kinases were defined according to the Illuminating the Druggable Genome program. Numbers within bars indicate kinase counts. Statistical comparison was performed using one-sided Fisher's exact test. ****$p$ < 0.0001.

**b,** Distribution of the 222 benchmark samples across perturbation classes. The ten focal perturbation classes used for class-level analysis are shown separately, whereas the remaining classes are grouped as other low-frequency perturbations for visualization.

**c,** Distribution of the same-family proportion across newly scoreable kinases included in the activity-correlation analysis. For each newly scoreable kinase, the same-family proportion was defined as the fraction of kinases scoreable using curated KSRs with valid activity correlations that belonged to the same kinase family.
